# Pulsation waves along the *Ciona* heart tube reverse by bimodal rhythms expressed by a remote pair of pacemakers

**DOI:** 10.1101/2023.10.06.561153

**Authors:** Yuma Fujikake, Keita Fukuda, Katsuyoshi Matsushita, Yasushi Iwatani, Koichi Fujimoto, Atsuo S. Nishino

**Affiliations:** Department of Biology, Graduate School of Agriculture and Life Science, Hirosaki University, Hirosaki 036-8561, Japan; Department of Bioresources Science, United Graduate School of Agricultural Sciences, Iwate University, Hirosaki 036-8561, Japan; Department of Biological Sciences, Graduate School of Science, Osaka University, Toyonaka 560-0043, Japan; Department of Science and Technology, Graduate School of Science and Technology, Hirosaki University, Hirosaki 036-8561, Japan; Department of Robotics, Faculty of Engineering, Kindai University, Higashi-Hiroshima 739-2116, Japan; Program of Mathematical and Life Sciences, Graduate School of Integrated Sciences for Life, Hiroshima University, Higashi-Hiroshima 739-8526, Japan

**Keywords:** heart, pacemaker, rhythm, oscillator, tunicate, ascidian

## Abstract

The heart of ascidians, marine invertebrate chordates, exhibits a tubular structure, and heartbeats propagate from one end to the other. The direction of pulsation waves intermittently reverses in the heart of ascidians and their relatives; however, the underlying mechanisms remain unclear. We herein performed a series of experiments to characterize the pacemaker systems in isolated hearts and their fragments and applied a mathematical model to examine the conditions leading to heart reversals. The isolated heart of *Ciona* sufficiently performed heart reversals, and experimental bisections of isolated hearts revealed that independent pacemakers resided on each side and also that their beating frequencies periodically changed as they expressed bimodal rhythms. Only fragments including 5% or shorter terminal regions of the heart tube maintained autonomous pulsation rhythms, whereas other regions did not. Our mathematical model, based on FitzHugh-Nagumo equations applied to a one-dimensional alignment of cells, demonstrated that the difference between frequencies expressed by the two independent terminal pacemakers determined the direction of propagated waves. Changes in the statuses of the terminal pacemakers between the excitatory and oscillatory modes as well as in their endogenous oscillation frequencies were sufficient to lead to heart reversals. These results suggest that the directions of pulsation waves in the *Ciona* heart reverse according to the changing rhythms independently expressed by remotely coupled terminal pacemakers.

**Summary statement:** Pulsation waves traveling along the heart tube of the ascidian *Ciona* intermittently reverse because of autonomous and periodical changes in beating frequencies at a pair of terminal pacemakers.

## INTRODUCTION

The heart is an essential organ in animals over a certain size and its rhythmic beating signifies the living status. The heart circulates body fluids for the delivery of nutrients and the exclusion of wastes from every part of the body. This is also the case in ascidians, which are marine invertebrates that have an open circulatory system. However, in the heart of ascidians and their kin, the direction of heartbeat waves intermittently reverses (e.g., Krijgsman, 1956; Schutt, 2021). This peculiar characteristic of their heart was initially reported approximately 200 years ago (Kuhl and van Hasselt, 1822); however, the underlying mechanisms have yet to be elucidated (Davidson, 2007; Cain et al., 2020).

The heart of ascidians is in the epicardium, a body cavity located between the branchial basket and stomach. The heart itself comprises two monolayered cell sheets, called the myocardium and pericardium (Fig. 1). The myocardium is a simple valveless tube called a heart tube that is composed of thin myoepithelial cells containing striated myofibrils (Kalk, 1970; Oliphant and Cloney, 1972; Nunzi et al., 1979). The heart tube is V-shaped in *Ciona*, an ascidian model genus that is utilized in many aspects of biological research (Satoh, 2013). The heart tube is present in the pericardium, and they are connected via a seaming structure called a raphe (Fig. 1) (Anderson, 1968; Kalk, 1970; Davidson, 2007; Anderson and Christiaen, 2016).

**Fig. 1.**
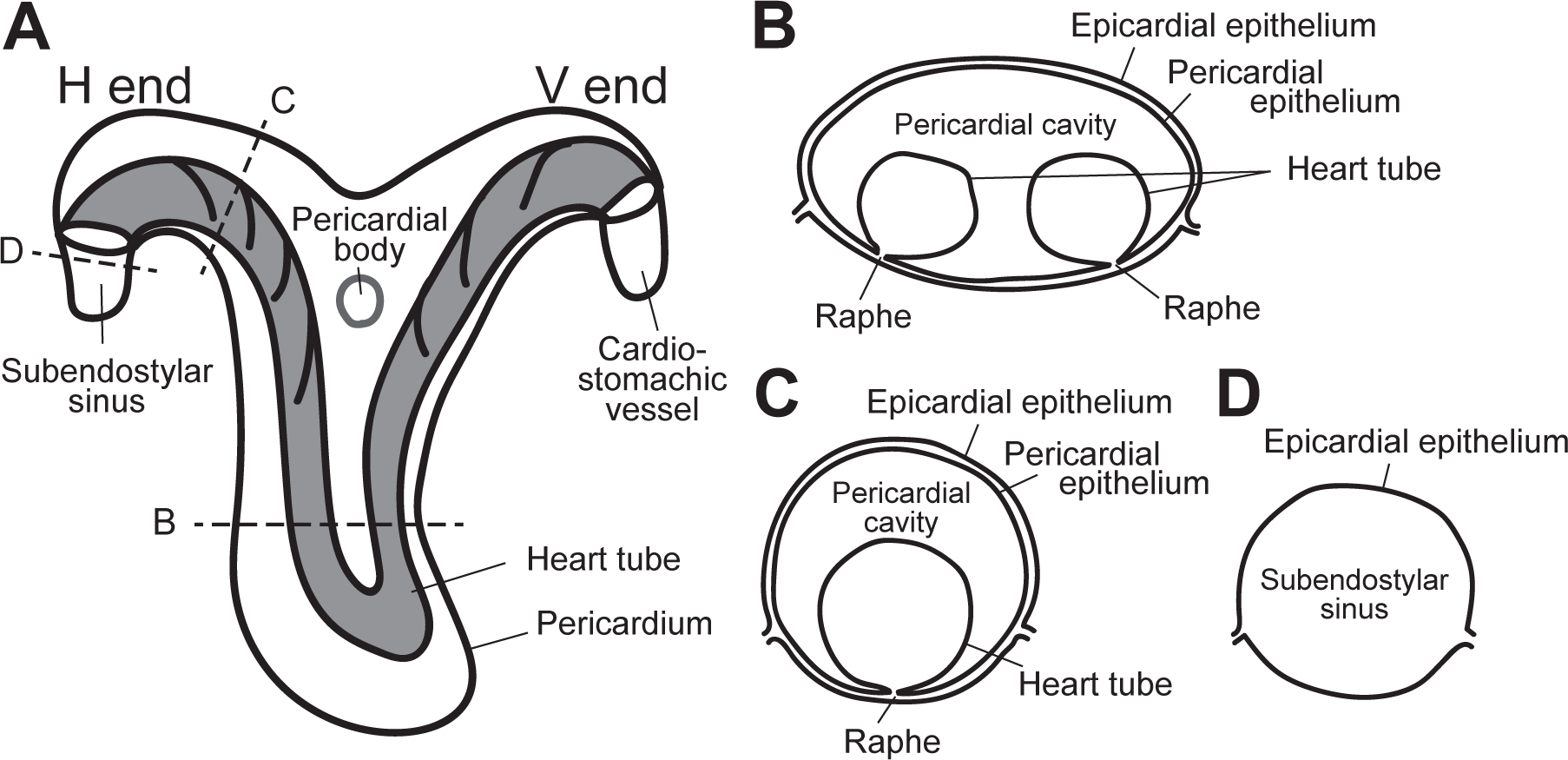
The structure of the *Ciona* heart. (A) A schema of the isolated heart. The heart tube (gray) has two ends, called the hypobranchial end (H end) and visceral end (V end), and is enclosed by the pericardium. (B-D) Cross-sectional images of the planes indicated in A. The heart tube is connected to the pericardial epithelium via the “raphe”, and the pericardium is covered by the epicardium.

One of the termini of the heart tube, the hypobranchial end (hereinafter called the H end), opens near the posterior end of the endostyle. The other end is adjacent to the stomach and, thus, is called the visceral end (hereinafter called the V end) (Kriebel, 1968a; Anderson, 1968). Both ends are connected to short vessels that soon open into the body cavity, thereby allowing body fluids and blood cells to move among tissues in an open circulatory system.

Pulsations in the ascidian heart tube appear to occur from one end to the other to elicit the directed flow of body fluids. However, the direction of pulsation waves intermittently reverses, and, accordingly, the direction of fluid flow also reverses. Why such reversals occur in ascidian hearts, i.e., ultimate causes for reversals, has not yet been established. Kuhl and van Hasselt (1822) speculated that circulating blood cells frequently clog narrow sinuses and, thus, the reversal may relieve this congestion (see Krijgsman, 1956; Heron, 1975). Subsequent studies attempted to connect this ultimate cause to a proximate cause, the so-called “back-pressure theory”. They predicted that the consecutive one-directional flow of blood cells may gradually increase resistance to circulation, which then triggers a reversal to obviate this elevation in pressure (Haywood and Moon, 1950; Kriebel, 1968a). However, this proposal was questioned based on the reports that periodic reversals occurred even in hearts isolated from the body (e.g. Millar, 1952; Krijgsman, 1956; Anderson, 1968). These reversals were also discussed to be related to some type of “fatigue” of the pacemaker(s) to generate heartbeats (Krijgsman, 1956).

The locations of pacemakers for heartbeats are also controversial. Since the pulsation waves propagated from either end of the tube, pacemakers were predicted to reside in terminal regions (Krijgsman, 1956). However, other regions have been suggested to autonomously contract to generate pulsation waves and, thus, pacemakers may be ubiquitously distributed along the heart tube (Anderson, 1968; Goodbody, 1974).

Ascidians are marine sessile invertebrates that represent the most closely related group to vertebrates. Based on this phylogenetic position, ascidians provide indispensable evidence for the origin and evolution of the vertebrate body. Despite the unusual nature of the ascidian heart, accumulating findings have revealed deep homology between gene regulatory networks for ascidian and vertebrate heart development (Satou et al., 2004; Davidson, 2007; Stolfi et al., 2010; Diogo et al., 2015; Wang et al., 2019; Razy-Krajka and Stolfi, 2019; Song et al., 2022). Under this context, questions regarding ascidian heart reversals are also expected to be re-interpreted in order to further refine our view of the phylogenetic establishment of the vertebrate heart (Davidson, 2007).

In the present study, we attempted to revisit this classic issue of ascidian heart reversals. We performed careful experiments on hearts in the heart in the body, on hearts isolated from the body, and on pieces of isolated hearts. We utilized modern techniques, such as video recording, motion analyses, and mathematical modeling. Our long-term recordings allowed us to estimate the extent to which artefactual effects affected the properties of the heart itself and its fragments. The results demonstrated that the heart tube isolated from the body sufficiently exhibited intermittent reversals. Further experimental fragmentations of isolated hearts showed that autonomous pulsations persisted in the fragments including either end with the property of bimodal rhythms. Our mathematical model demonstrated that independent changes in the rhythms expressed by a pair of remotely coupled pacemakers were sufficient to lead to heart reversals. These results suggest that an intrinsic property of the pair of terminal pacemakers to express independent bimodal rhythms leads to heart reversals.

## MATERIALS AND METHODS

### Animals

The hearts of adult *Ciona robusta* Hoshino and Tokioka, 1967, also called *C. intestinalis* (Linnaeus, 1767) type A, were used in experiments in the present study. Adult animals were obtained from the Misaki Marine Biological Station, The University of Tokyo (Miura, Japan), or the Maizuru Fisheries Research Station, Kyoto University (Maizuru, Japan) under the support of the National BioResource Project, AMED, Japan. The adult individuals delivered were kept in the laboratory as previously described (Jokura et al., 2020; Hara et al., 2022). We performed the following experiments along the procedures in compliance with the animal welfare guidelines defined by the affiliation (Hirosaki University).

### Preparation of hearts

All experiments were performed in an air-conditioned room (18-21). *Ciona* adults were placed in a Petri dish containing artificial seawater (ASW; Marine Art BR, Osaka Yakken, Japan). The tunic was cut open and removed with scissors, the posteroventral part of the body wall (the mantle) was then opened with tweezers, and the heart was carefully exposed through this opening.

To isolate the heart, the two connections of the heart tube to other parts of the body, e.g. the subendostylar sinus on the side of the H end and the cardio-stomachic vessel on the side of the V end, as well as other connections of the heart if necessary, were cut off using micro-scissors (vannas scissors, type 14003, World Precision Instruments). The isolated heart was placed in another Petri dish containing ASW, the bottom of which was covered with silicone rubber (SYLGARD 184, Dow Corning). Bisections and further divisions of the heart tube were performed in the silicone-covered Petri dish containing ASW.

### Acquisition of video images

Pulsation waves generated in heart tubes were recorded with a video camera (HDR-CX420, SONY) mounted on a stereomicroscope (SMZ745T, Nikon). Pulsation waves in or out of the body were recorded.

Regarding the recording of heartbeats before isolation (see Results), the adult body from which the tunic had been removed or, in some cases, in which the heart was partially exposed was laid in ASW on the silicone rubber pad in the deep Petri dish. Body-wall muscles that were distant from the heart were pinned down on the silicone pad with cactus spines, which prevented overall shrinkage of the body. Immediately after the operations, spatiotemporal patterns of pulsations of the heart viewed through the translucent mantle or those of the exposed heart were recorded.

To observe isolated hearts, specimens were placed in ASW on the silicone rubber pad in a 35-mm Petri dish. Several points along the outline of the epicardial and pericardial epithelial sheets were carefully pinned down using cactus spines. The pulsations of the heart were recorded immediately after the isolation as above. The conditions for recording pulsation patterns in bisected or divided heart fragments were similar, and rest periods in the dish were as denoted in the Results. Movie files for presentations were prepared with the video editor Clipchamp running on a Windows PC.

### Image analysis

Recorded movie files [29.97 frames sec^-1^ (fps), mts or m2ts format] were exported into an image sequence of TIFF- or BMP-formatted images at 10 fps using a locally developed application running on a Windows PC (Hara et al., 2022). We approximated the original video as 30 fps, and the resulting error of one frame per 1000 s was ignored. The exported image sequence was imported into ImageJ (version 1.48, NIH, USA). Changes in 8-bit-scaled brightness (0-255) at a “point of interest (POI)” were quantified and depicted. When the shade or reflection on the heart tube traversed the POI as many times as the heart pulsated, brightness at the POI changed (decreases or increases, respectively) with heartbeats.

Time series data on quantified brightness at the POI was imported into Excel (Microsoft), and the frequency of heartbeats (beats per minute, bpm) was calculated by dividing the time duration of multiple beating cycles by the number of cycles (ranging from 2 to 35 cycles, depending on the conditions). In some cases, the durations of beating cycles were directly measured by observing the movies using a stopwatch.

After the detection of heart reversals in the movie by the naked eye, the durations of the H- or V-dominant series of contraction waves were measured. Heart reversal did not occur in one out of 13 examples or only occurred once during recordings for 2 h or longer. These two examples were excluded from quantitative analyses (see the legend for Fig. 2). Averaged values for the duration of H- or V-dominant periods in 1-h movies were treated as the representative value of the heart tube. We sometimes (or often after isolation) observed that the hitherto dominant side remained dominant even after a series of collisions. These cases were regarded as the dominant period of the hitherto dominant side continuing even during collisions.

**Fig. 2.**
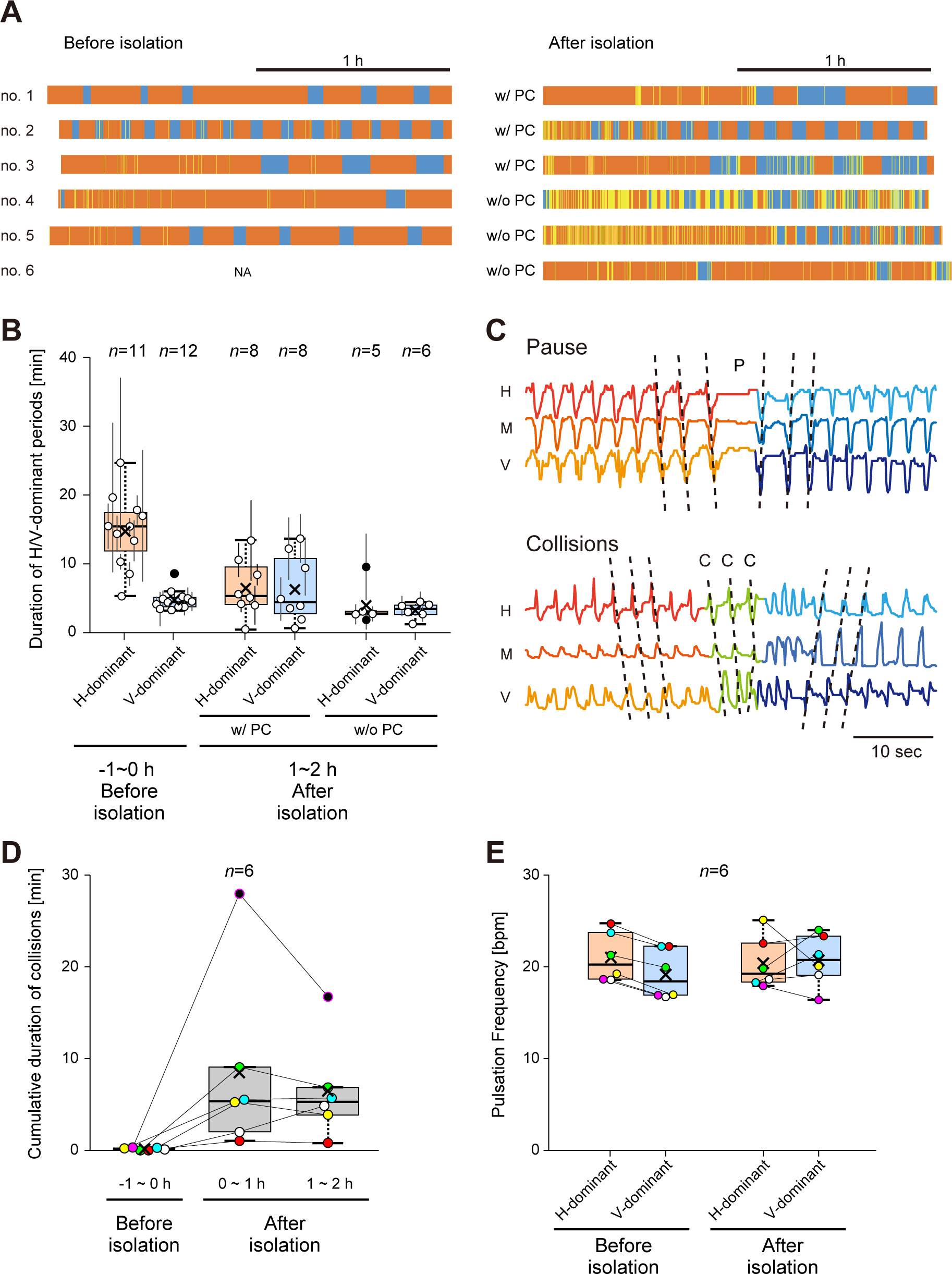
Patterns of pulsation waves in *Ciona* hearts before and after isolation. (A) The pulsation patterns of five independent hearts viewed through the mantle or partially exposed through a nick on the mantle (left), and those of the identical hearts after isolation (right). Tangerine, cyan, and yellow bars indicate the H- and V-dominant periods, and the periods in which collisions occurred, respectively. The pericardial wall was intact in specimen no. 1–3 (w/ PC), whereas that was cut open in no. 4–6 (w/o PC). (B) Duration of H- and V-dominant periods in the heart before isolation (−1∼0 h), 0∼1 h after isolation, and 1∼2 h after isolation. Plots and vertical bars indicate the mean duration and its standard deviation in each heart, respectively. Two samples were excluded from the analysis, since no or only one reversal occurred during recordings for 2 h. (C) The heart reversal with a pause and collisions. Pulsation waves were initially propagated from H to V in the indicated cases. In the “pause” (upper), the pulsation ceased for a while (indicated with “P”), and another series of pulsations were then propagated from V to H. In the “collisions” (lower), pulsation waves from both ends collided on the way (labeled with “C”) and pulsations then propagated from V to H. (D) Cumulative durations of collisions in the heart 1∼0 h before isolation (−1∼0 h), 0∼1 h after isolation, and 1∼2 h after isolation. Plots with the same colors indicate data from identical hearts. (E) Pulsation frequencies during the H- and V-dominant periods in hearts before and after their isolation. Data from identical hearts were represented by the same colors and connected with colored lines. For the panels B, D, and E, × and *n* indicate the mean value and the number of examined hearts in each condition, respectively. Plots with a filled black color are outliers.

### Beating frequency analysis through wavelet transformation

Recorded movie files were transformed into 10 fps movies using FFmpeg (https://www.ffmpeg.org/). The averaged background image was subtracted from every frame and also transformed into 8-bit grey-scale AVI movies using OpenCV (version 4.5.3). These movies were opened using ImageJ, and changes in the brightness value, which was averaged in regions of interest (the ROI) that covered the H or V end, were quantified and exported as time-series data. We normalized changes in values during periods of interest to a range between 0 and 16, because changes in brightness values were too small in some cases for further analyses. Using the PyWavelets library which ran on Python3, time-series data were examined through a continuous wavelet transformation, and the power spectrum of every frequency was calculated along the time course and depicted by a heat map. Continuous wavelet transformation is represented by the following equation:

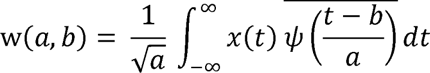

where *x* represents a time-series signal. The parameters *a* and *b* are involved in the scaling and time shift, respectively, of the mother wavelet ψ, where 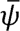, denotes the complex conjugate. In the present study, the mother wavelet was defined as a complex Morlet wavelet function as follows:

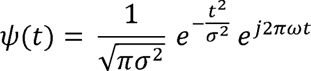

In the present study, we set *σ*^2^ and ω at 1.5 and 1, respectively. The value *j* represents the imaginary unit. In this analysis, frequencies were examined by 0.001 Hz increments in the range of 0.10–1.0 Hz (6.0–60 bpm). The resolution of time on the heat maps was defined as 0.10 s. Frequencies lower than 0.10 Hz (= 6.0 bpm) were ignored and represented as 0.10 Hz (= 6.0 bpm), and, thus, the peak frequency of the wavelet power spectrum was taken into consideration after a cut-off < 0.12 Hz using fourth-order high-pass filtering in Python. The time-series data obtained on peak frequency were plotted along the temporal axis and compared with the reciprocal numbers of the intervals of pulsation events. The intervals of the pulsation events were measured as peak-to-peak durations in the time-series data derived from the brightness changes on the image sequences.

### Mathematical modeling

In the mathematical model of pulsation waves observed in the *Ciona* heart tube, we made the following assumptions. Pulsation waves propagating along the heart tube were a representation of action potential (AP) propagations. All of the cardiac cells in the heart tube are excitable and are connected via gap junctions (Kalk, 1970; Nunzi et al., 1979; Anderson, 1968). In addition, based on the present results (see Results), we assumed that pacemaker cells that autonomously generated APs were located in the terminal regions of the heart tube, and working myocytes that were passively excited in response to the depolarization signal propagating from neighboring cells were located between pacemaker cells. We simply modelled the *Ciona* heart tube as a one-dimensional multicellular tissue composed of 20 equal-sized cells, in which single pacemaker cells (hereinafter referred to as P cells or P_H_ and P_V_ cells) were placed at each of the ends and the remaining 18 working cardiomyocytes (W cells) were linearly aligned between P cells. This configuration reflected the present results showing that the regions of pacemakers were restricted to 5% or shorter terminal regions (see Results).

We utilized FitzHugh-Nagumo equations to generate APs in our model (FitzHugh, 1961; Nagumo et al., 1962). FitzHugh-Nagumo equations have been proposed as a versatile mathematical model that simplifies Hodgkin-Huxley equations (Hodgkin and Huxley, 1952). Our model is represented by the following equations:

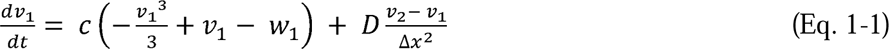

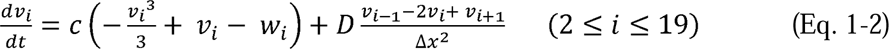

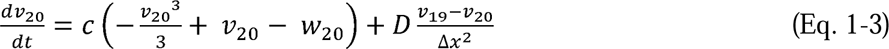

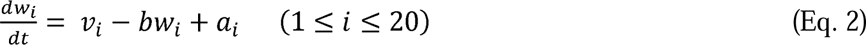

in which *v* and *w* represent the membrane potential and open probability of ion channels, respectively, in cell “*i*” (1 ≤ *i* ≤ 20). Eq. 1-1 and 1-3 represent P cells with the H- and V-marginal status, respectively, reflecting the condition that cell “1” and cell “20” receive/pass the signal only from/to cell “2” and cell “19”, respectively, unless otherwise specified. On the other hand, Eq. 1-2 represents W cells, where cell “*i*” receives/passes the signal from/to cells “*i*-1” as well as “*i*+1”. Parameters “*a*” and “*b*” are involved in restricting the excitability as well as the ability for spontaneous oscillations in each cell. The decrease in “*a*” increases the excitability and a further reduction elevates the autonomy of its excitation. “*b*” and “*c*” are parameters that change the relative excitability of cells, but were commonly fixed in all cells in the present study, whereas the values for “*a*” were independently defined among cells; therefore, it was depicted as “*a_i_*” (1 ≤ *i* ≤ 20). “*D*” and “*Δx*” are diffusion constants, reflecting the relative conductibility of APs and the size of the cells, respectively.

The FitzHugh-Nagumo model allows us to describe two modes; APs are autonomously and repetitively generated in one mode (the oscillatory mode), while Aps are passively generated in response to depolarizing signals propagated from neighboring cells in the other mode (the excitatory mode). Continuous changes in the values of parameter “*a*” (or those of “*b*” or “*c*”) lead to discontinuous transitions between the oscillatory and excitatory modes, or a frequency change in the AP oscillation within the oscillatory mode (see also Results). In the present study, the parameters “*a_i_* (1 ≤ *i* ≤ 20)” were set to characterize P cells as oscillatory cells and W cells as excitatory cells.

Eq. 1 and 2 were numerically calculated using the Runge-Kutta method (fourth order) written in C language. The temporal increment and other parameters were defined as *dt* = 1.0 × 10^-3^, *b* = 0.80, *c* = 10, *D* = 5.0 × 10^-6^, and *Δx* = 1.0 × 10^-3^, respectively. The initial values of P cells (*i* = 1 for P_H_ and *i* = 20 for P_V_) and W cells (2 ≤ *i* ≤ 19) were also defined as (*v_i_* = −5.0 and *w_i_*= 0) and (*v_i_* = −8.0 and *w_i_* = 0), respectively. Subsequent changes in the membrane potential (*v*) and open probability of ion channels (*w*) were computed for all cells. The results obtained were visualized using matplotlib of Python3.

## RESULTS

### Heart reversals in *Ciona* occur in an autonomous manner

We examined pulsation waves occurring in the heart of the adult body. We stripped the tunic of adult *Ciona* and, in some cases, made a nick on the mantle to partially expose the heart. We then immediately recorded the patterns of pulsation waves on the heart tube for 1 h or longer. The duration of the H-dominant period, in which repetitive contraction waves were propagated from the H end to the V end, was longer and varied more than that of the V-dominant period (Fig. 2A,B). The duration of the V-dominant period was short and constant (3–9 min) (Fig. 2A,B).

We then monitored the patterns of pulsation waves in hearts outside of the body (Fig. 2). In some cases, we also cut open the epicardial and pericardial epithelia enveloping the heart tube (Fig. 2A,B). Immediately after isolation, a long H-dominant period persisted for tens of minutes to 1 h, whereas heart reversals revived 1 to 2 h after isolation or later (Fig. 2A). Importantly, two independent patterns of heart reversals, i.e., a pause or a series of collisions, were also observed in isolated hearts, similar to hearts within adult bodies (Fig. 2C) (Movie 1) (Millar, 1952; Krijgsman, 1956). In the former cases, pulsation waves from one end ceased, a short pause followed, and then those from the opposite end began (Fig. 2C, top). In the latter cases, before the reversal was completed, pulsation waves travelling from both ends collided midway on the heart tube for several cycles (Fig. 2C, bottom) (Movie 1).

Although heart reversals were qualitatively restored in isolated hearts, significant quantitative changes were detected in some of the temporal parameters. The duration of the H-dominant period was significantly shorter after than before isolation (p < 0.01, the Student’s *t*-test, two-tailed, the same applies below unless otherwise specified), whereas changes in the duration of the V-dominant period were not apparent (p > 0.8) (Fig. 2A,B). Also, collisions were frequently seen in the isolated hearts (Fig. 2A,D).

To further estimate the effects of isolation, pulsation frequencies were compared before and after isolation. In hearts before isolation, pulsation frequencies recorded during the H-dominant period were higher than those during the V-dominant period in all cases (Fig. 2E). On the other hand, in hearts 1 h or more after isolation, this asymmetry disappeared and pulsation frequencies in the H-dominant period were not always higher than those during the V-dominant period (Fig. 2E). These changes after isolation suggested that the surgical manipulations applied to the hearts exerted some artefactual effects on the pulsation pattern.

We examined the pulsation frequencies in hearts from immediately to 1 day after their isolation. The results obtained demonstrated that pulsation frequencies significantly decreased from 30.7 ± 1.8 bpm (mean ± standard deviation; n = 10) immediately (1–4 min) after isolation to 23.2 ± 1.6 bpm (n = 10) 2–4 h after isolation (p < 0.01) (Fig. 3A,B). Isolated hearts did not exhibit further notable changes even after 1 day (23.5 ± 2.8 bpm, n = 6; p > 0.5, vs. that 2–4 h after isolation) (Fig. 3A,B). Intermittent heart reversals were also observed even one day after isolation (Movie 2).

**Fig. 3.**
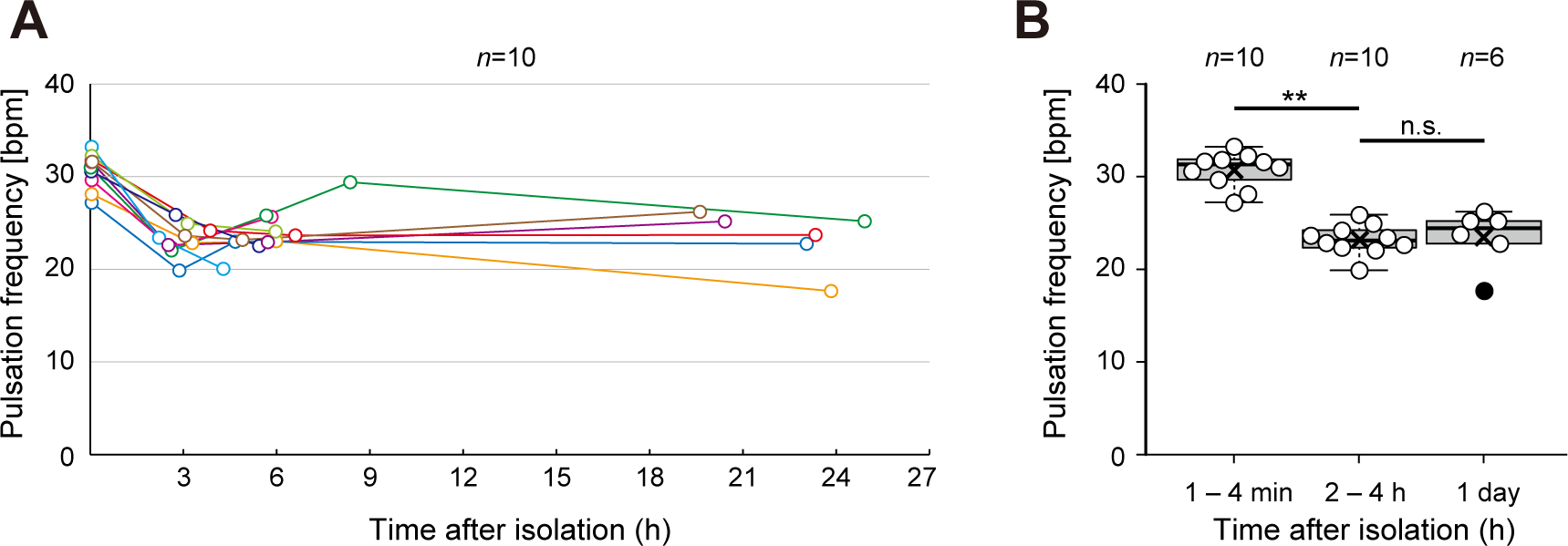
Changes in pulsation frequencies after isolation. (A) The pulsation frequencies of isolated hearts immediately, 2–4 h, 4–9 h (in some examples), and 1 d after isolation. Data from identical hearts were connected with colored lines. (B) Boxplots of pulsation frequencies immediately (1–4 min), 2–4 h, and 1 d after isolation. × and *n* indicate the mean value and the number of data in each condition. ** and n.s. indicate p < 0.01 and p > 0.05 (not significant), respectively (the Student’s *t*-test, two-tailed).

Although some changes were detected in hearts after their isolation, pulsation waves and their reversals continued for one day or longer. These results allowed us to estimate artificial effects in the operated hearts, and also informed us that heart reversals in *Ciona* occurred sufficiently in an autonomous manner. We considered examinations of heart reversals in isolated hearts to be still important for elucidating the intrinsic mechanisms responsible for heart reversals in *Ciona*, and, thus, further investigated pulsation patterns in isolated hearts or their fragments as described below.

### Changes in beating rhythms are associated with heart reversals

We analyzed changes in pulsation frequencies at the H and V ends immediately before and after heart reversals in isolated hearts. We examined five cases of heart reversals with a pause. Wavelet transformation was applied to analyze time-series data with a high temporal resolution (see Methods), and the results obtained revealed that pulsation frequencies gradually decreased before the pause occurred (Fig. 4A; 100%, n = 5).

**Fig. 4.**
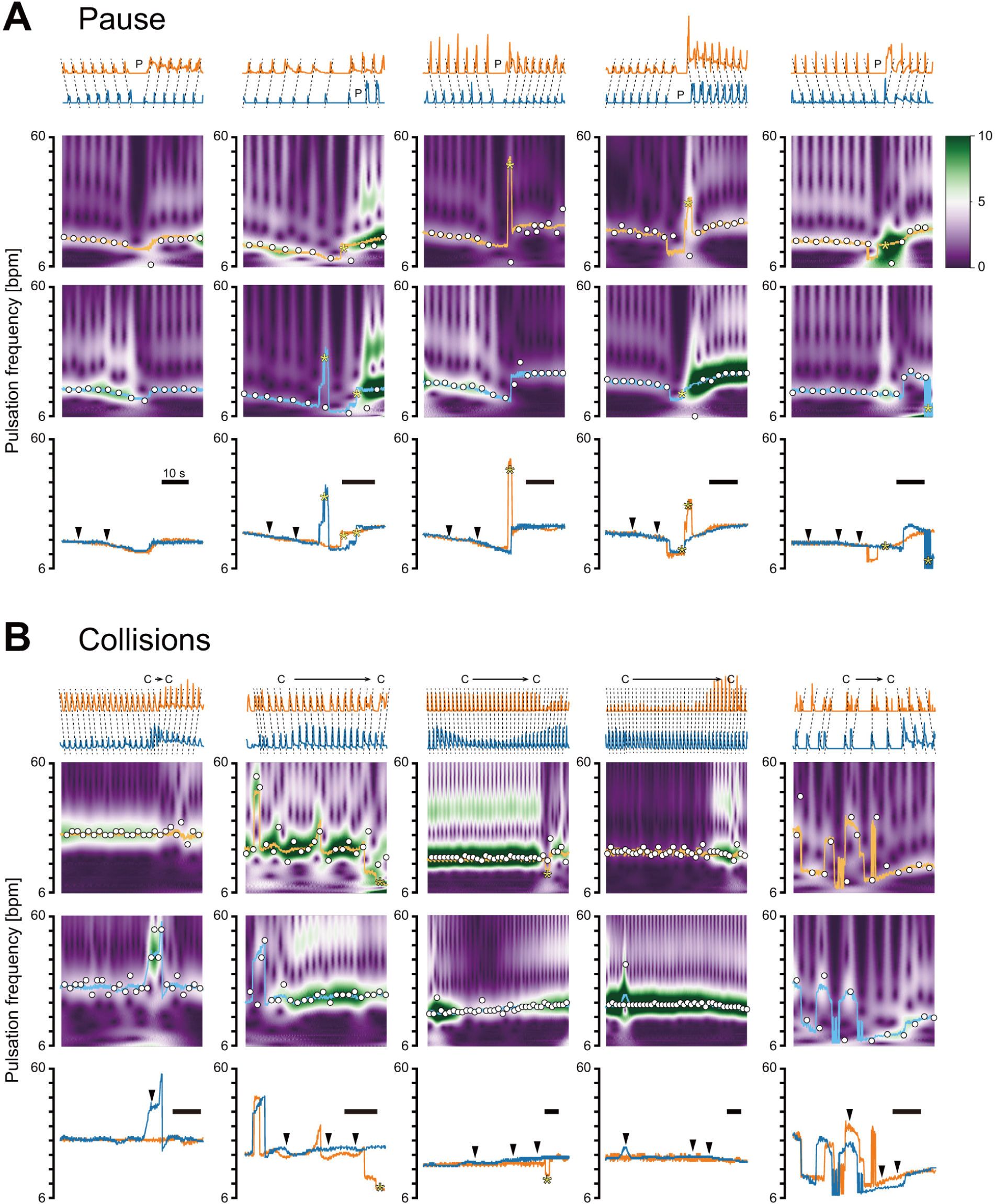
Changes in pulsation frequencies around heart reversals. (A) The pulsation frequency gradually decreases before heart reversals with a pause occur. (B) Pulsation frequencies in the hitherto recessive end overcome those in the hitherto dominant end when heart reversals with collisions occur. The top traces indicate the beating pattern in the H- (tangerine) and V- (cyan) ends, representing brightness changes in ROIs including the ends (arbitrarily scaled). Dashed lines show the wave direction (H to V, or V to H). “P” and “C” mark the timing of pauses and collisions, respectively. The middle two rows are heat maps showing the wavelet power spectra in the H (upper) and V (lower) ends, respectively. The peak frequencies of the high-pass filtered (> 0.12 Hz) wavelet power spectrum at each time point are plotted with light tangerine (H) and cyan (V) lines. The reciprocal values of beating intervals are also merged (open circles). These plots appropriately fit the wavelet power spectrum. Acute changes in the peak frequency that did not appear to reflect the beating pattern are marked with yellow asterisks. Traces of the peak frequencies of wavelet power spectra in the H and V ends are compared at the bottom. Arrowheads indicate gradual decreases (A) or overcoming of the frequency by the hitherto recessive end (B). Bars = 10 s. The other two examples are shown in Fig. S1.

While collisions occurred, pulsation frequencies independently recorded at the terminal regions were assumed to be the intrinsic rhythm expressed by each end. We examined seven cases of heart reversals with collisions and noted that the pulsation frequency in the hitherto recessive end tended to gradually or sometimes acutely overcome that in the hitherto dominant end before the reversal was accomplished (Fig. 4B and Fig. S1; 100%, n = 7). These results suggested that the pulsation rhythms appearing in the H and V ends could change and the occurrence of heart reversals was related to changes in the rhythms expressed in the ends.

### The pulsation rhythm in the dominant end persists after bisection of the heart tube

In order to know what would happen in the recessive end if the travelling waves from the other end would not come there repeatedly, we divided the heart tube into the H and V halves. Immediately after bisection of the heart tube, pulsation waves were generated not only from the dominant end but also from the hitherto recessive end to the cut site (Movie 3).

We compared wavelet-transformed pulsation frequencies for 40 s before and 40s after the bisection. When the heart tube was divided during the H-dominant period, the pulsation frequency was maintained in the H end after the bisection (100%, n = 6), and the frequency in the H end was higher than that in the V end after the bisection (100%, n = 8) (Fig. 5A and Fig. S2A). Moreover, when the heart tube was divided during the V-dominant period, the pulsation frequency was more likely to persist in the V half after the bisection (87.5%, n = 8) (Fig. 5B and Fig. S2B), except in one case in which the V and H halves paused immediately after the bisection (Fig. S2C). In the examples in which beating persisted in the hitherto dominant end, the pulsation frequency in the V end was higher than that in the H end after the bisection (71.4%, n = 7) (Fig. 5B and Fig. S2B). These results indicate that the pulsation frequency in the dominant end was maintained in the hitherto dominant half even after the bisection (93.8%, n = 16 in all), and the frequency observed in the hitherto recessive side after the bisection was lower than that in the hitherto dominant side (86.7%, n = 15 in all).

**Fig. 5.**
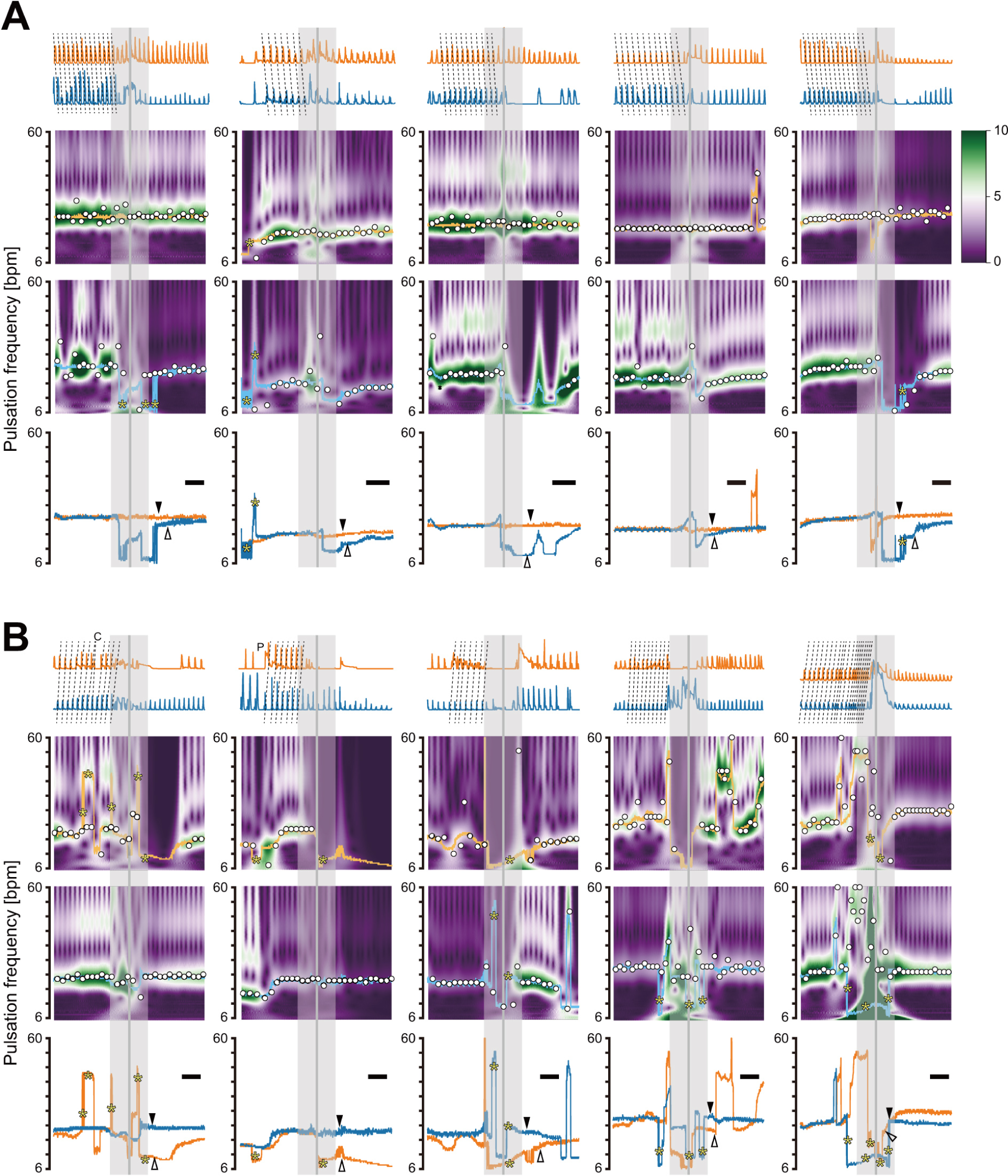
Changes in pulsation frequencies before and after bisection of the heart tube. (A,B) Changes in pulsation frequencies in the H (tangerine) and V (cyan) ends before and after the bisection in the H-dominant (A) and V-dominant (B) periods. The indications are the same as in Fig. 4. Vertical lines and grey boxes mark the timing of the bisection and ± 10 s range before and after the bisection. Black and white arrowheads indicate the peak frequencies of the power spectra in the hitherto dominant and recessive ends, respectively. Bars = 10 s. Other examples are shown in Fig. S2.

### Bimodal rhythms expressed by H and V ends

To investigate the nature of each of the halves, the beating patterns of the H and V halves were recorded for 30 min immediately after and 1.5 h after the bisection (n = 7) (Fig. 6A,B). During the first 30 min after the bisection, pulsations in the V halves sometimes paused (57.1%, n = 7), whereas those in the H halves remained regular (100%, n = 7) (Fig. 6A).

**Fig. 6.**
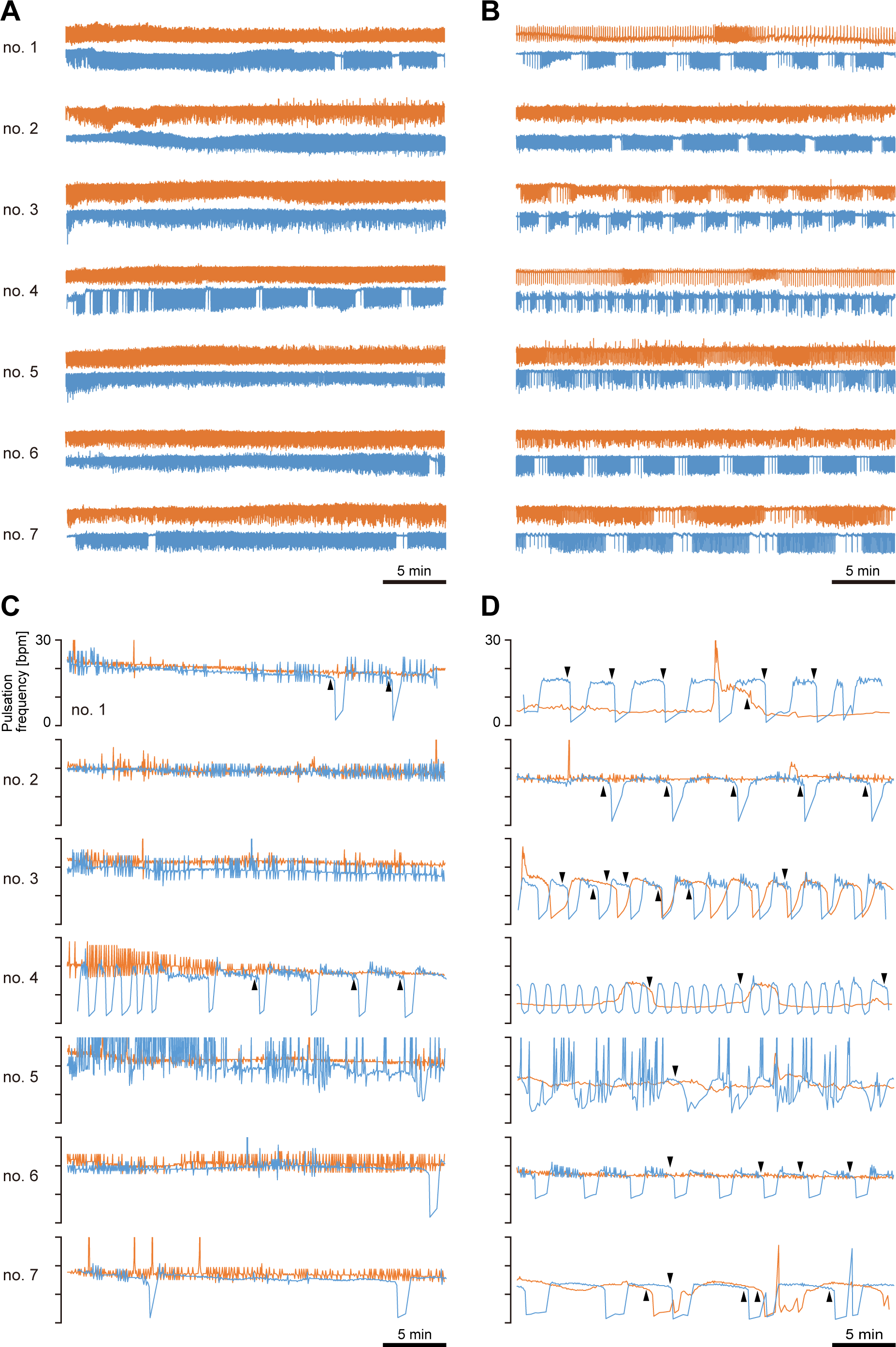
Periodic changes in pulsation frequencies after bisection of isolated hearts. (A) Pulsation patterns of the H and V halves for 30 min immediately after (A) and 1.5 h after the bisection (B). (C,D) Temporal changes in the pulsation frequencies, represented by the reciprocal values of the beating intervals. Frequency changes appeared to be associated with a gradual and acute phase. Gradual decreases before acute declines are indicated with arrowheads.

In the 30 min that followed 1.5 h after the bisection, repetitive accelerations or decelerations of the beating frequency were evident in the V halves (100%, n = 7), and similar temporal patterns were also noted in some of the H halves (71.4%, n = 7) (Fig. 6B). Time-series data represented by reciprocals of the beating intervals showed that decelerations were very rapid; however, slow decreases in the frequency were often observed before these acute declines (Fig. 6C,D; arrowheads).

These repetitive changes in pulsation frequencies appeared to have a cycle of approximately 1.25–5.5 min (= 0.003–0.013 Hz) (Fig. 6B,D). This result implied that the H and V halves both possessed an intrinsic property to generate compound rhythms of ∼0.003–0.013 Hz and ∼0.25–0.42 Hz (= ∼15–25 bpm). The properties on both sides of the heart tube to express bimodal rhythms, namely, a ∼1.25–5.5-min acceleration/deceleration cycle of a beating rate with a maximum of ∼15–25 bpm, support the results described above on the pulsation rhythms generated by the H and V sides changing by their own intrinsic mechanism.

### Pacemakers of the *Ciona* heart are restricted within terminal regions

To localize the pacemaker regions, we divided heart tubes into quarters and recorded their beating patterns (Fig. 7, Fig. S3). In the fragments including the H or V end (the H or V terminal quarters), pulsations persisted for 30 min or longer (100%, n = 10) (Fig. 7A, Fig. S3). The H and V terminal quarters sometimes exhibited pauses or acceleration-deceleration cycles during the first 30 min after divisions (50% and 90% of the H and V terminal quarters, respectively, n = 10 in each); however, pulsations were maintained in all cases (100%, n = 20 in total) (Fig. 7A, Fig. S3). Several quarters excluding the H and V ends (the H or V middle quarters) also showed rhythmic beating during the first 10 min after divisions (70% and 30% of the H and V middle quarters, respectively; n =10 in each). However, in all observed cases, beating frequencies in the middle quarters were lower than those in the terminal counterparts, and beating in the middle quarters gradually slowed and finally disappeared within 15 min (100%, n = 20 in total) (Fig. 7A, Fig. S3).

**Fig. 7.**
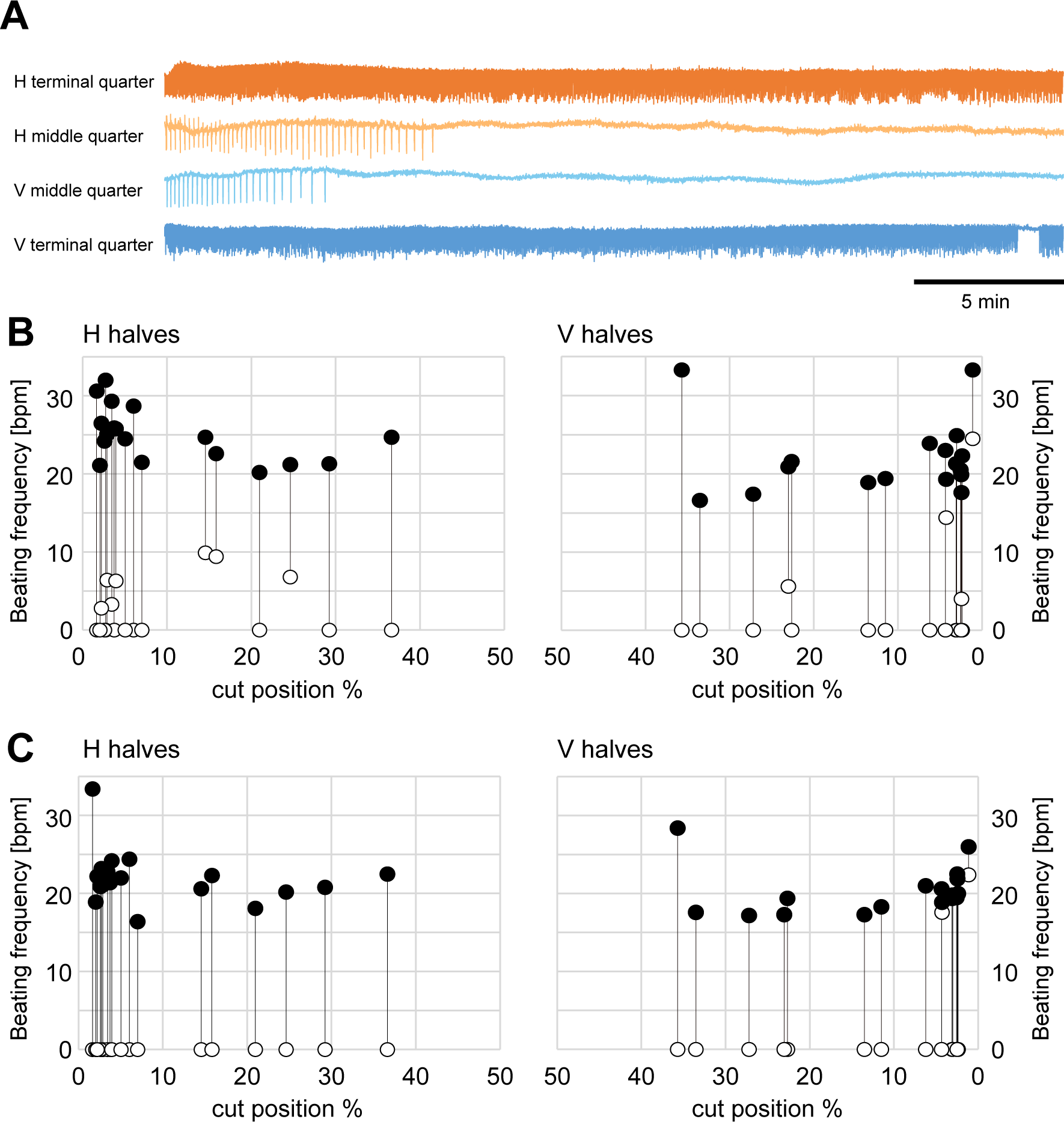
Fragments including terminal regions sustained autonomous pulsations. (A) Isolated hearts were bisected into the H and V halves, which were then subdivided into the terminal and middle quarters, respectively. Representative beating patterns in these quarters are presented here, and other examples are shown in Fig. S3. Tangerine and cyan traces show the beating patterns of the H- and V-sided terminal quarters, while light tangerine and cyan traces show those of the H- and V-sided middle quarters, respectively. (B,C) Isolated hearts were cut at the 50% position (when the length of the whole heart tube was regarded as 100%), and the H and V halves were then subdivided at arbitrary positions. The beating frequencies of the fragments immediately after divisions (B) and 30 min after divisions (C) were analyzed during the regular beating phase, and plotted against the positions of the second cut (cut position). Pairs of filled and open circles (connected with vertical lines) indicate the beating frequencies of the terminal and middle fragments, respectively, derived from identical halves.

We examined the potential of terminal fragments to sustain beating. We divided the heart tube into halves, and then cut each once more at an arbitrary position (Fig. 7B). We recorded the beating patterns of the terminal and middle fragments and examined the relationship between the cut positions and beating frequencies of the fragments (Fig. 7B,C). In the period immediately after the divisions, not only all of the terminal fragments (100%, n = 35), but also some of the middle fragments (31.4%, n = 35) exhibited rhythmic beating (Fig. 7B, filled and open circles, respectively). When the middle fragments showed beating, their frequencies were lower than those of the terminal counterparts (100%, n = 11 in total) (Fig. 7B).

Approximately 30 min after the divisions, all of the terminal fragments sustained beating (Fig. 7C). In most of the terminal fragments (88.6%, n = 35), beating frequencies became lower than those immediately after the divisions (23.5 ± 4.3 vs. 21.2 ± 3.3 bpm, n = 35 in each; p < 0.05), implying a decline in the artefactual effect (Fig. 7B,C). Short fragments (length < 5% of the whole heart tube) sufficiently had the potential to express bimodal rhythms 30 min after the division (Movie 4).

On the other hand, most of the middle fragments ceased beating by 30 min after the divisions (Fig. 7C). There were two exceptions, in which V middle fragments cut at 4.3% and 1.1%, respectively, sustained beating (n = 35) (Fig. 7C). These results suggested that pacemakers of the *Ciona* heart resided in terminal 4.3% or shorter regions.

### Differences in the frequency of remotely coupled pacemakers determine the direction of propagating APs

To build our mathematical model of the heart tube based on FitzHugh-Nagumo equations (Eqs. 1 and 2), we defined a cell population and investigated the parameter conditions evoking wave propagations over the cells. A single pacemaker cell (P cell) was placed on one of the ends and multiple working myocytes (W cells) were linearly aligned beside it (Fig. 8A). We initially set the model parameters of W cells to be in the excitatory (passive) mode but gave specific parameters that were close to the threshold for expressing autonomous oscillations (specifically, *a* = 0.44, *b* = 0.80, *c* = 10; see Methods). Under this condition, when the control parameter “*a*” in the P cell (*a*_1_ = 0.40) was below the threshold, the P cell oscillated, and APs were periodically propagated (Fig. 8B). The frequency of propagated APs increased as “*a*” in the P cell decreased (Fig. 8C), corresponding well with the definition that “*a*” is a parameter to restrict the excitability. Therefore, we fixed the parameter of the P cell at this level (*a*_1_ = 0.40) and also fixed that of W cells in the excitatory mode (*a* ≥ 0.41). Model simulations also showed that above another threshold of the parameter (*a* ≥ 0.45) in W cells, APs generated in the P cell did not travel over W cells (Fig. 8D). This result suggests that W cells need to possess excitability above a certain level close to that in the oscillatory pacemaker (0.45 > *a* ≥ 0.41). Based on this result, we set the parameter “*a*” of W cells slightly above the threshold of the oscillatory mode (*a* = 0.44) in subsequent experiments.

**Fig. 8.**
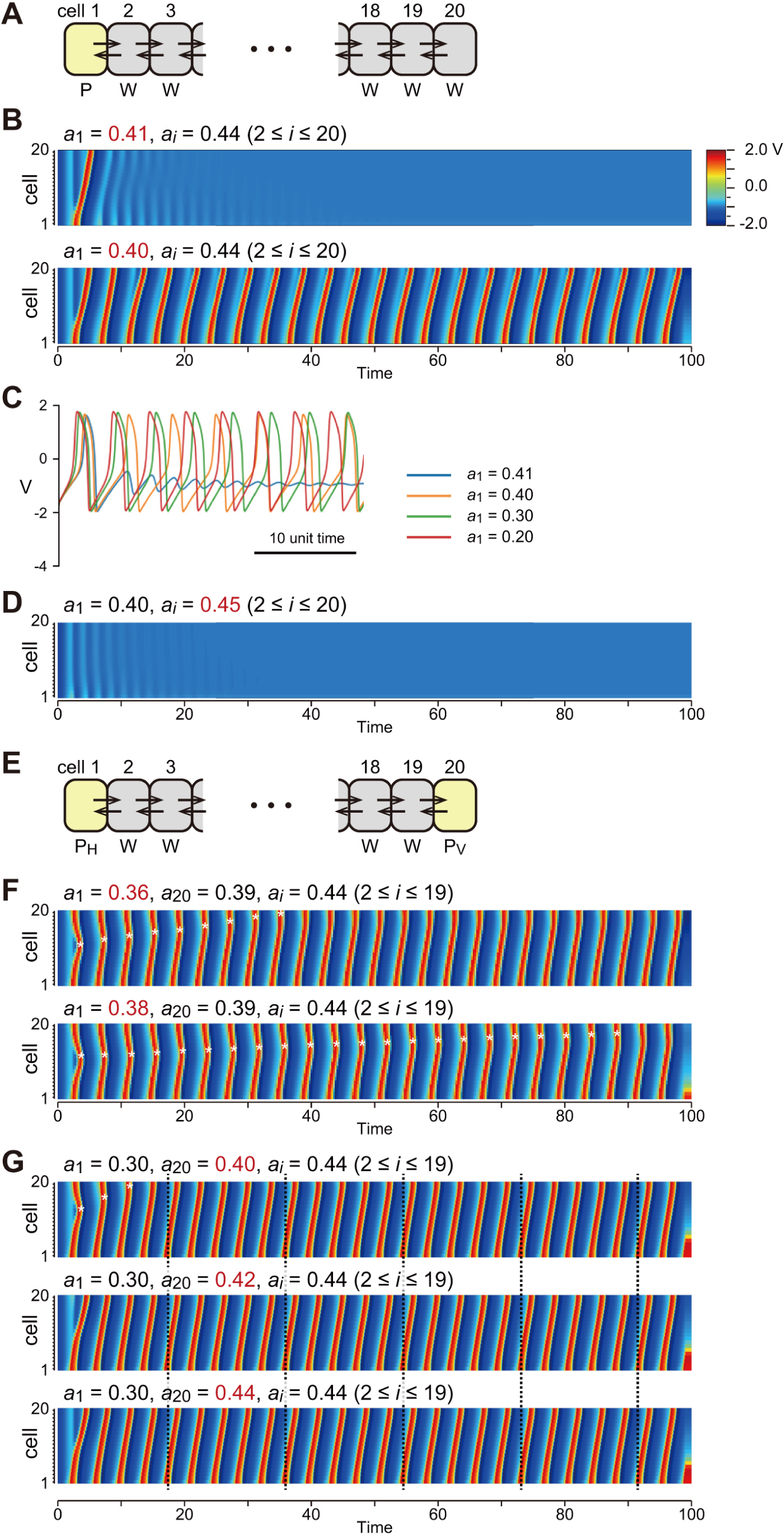
Preparation of a *Ciona* heart tube model. (A) Arrangement of the model cells. A pacemaker cell (oscillatory mode, labeled with “P”) was placed at the end and the other 19 cells (excitatory mode, “W”) were linearly aligned. (B) When parameter “*a*” in the P cell was above the threshold (*a*_1_ = 0.41; excitatory mode), pulsation waves did not repeat (upper). When parameter *a*_1_ was below the threshold (*a*_1_ = 0.40; oscillatory mode), pulsation waves were rhythmically propagated. (C) Decreases in parameter *a*_1_ increased the frequency of pulsation waves. (D) Even if the P cell was in the oscillatory mode (*a*_1_ = 0.40), W cells with less excitablity (*a_j_* = 0.45, 2 ≤ *j* ≤20) did not support the wave propagation. (E) Arrangement of the “two-pacemakers” model. A pair of P cells (P_H_ and P_V_) were placed at the ends, and W cells were aligned in between. (F) P_H_ and P_V_ both generated pulsation waves and collisions occurred, when they possessed a sufficiently low “*a*” (*a*_1_ = 0.36 for the P_H_, *a*_20_ = 0.39 for the P_V_) (upper). When *a*’s in P_H_ and P_V_ were closer, the duration of collisions was extended (lower, cf. upper). (G) When one of the P cells dominated to generate pulsation waves (*a*_1_ = 0.30), changes in “*a*” in the other (*a*_20_ = 0.40, 0.42, and 0.44) did not give rise to any differences in the pulsation pattern after the period of collisions ended.

We placed two P cells (P_H_ and P_V_) on both ends to examine what determines the direction of AP waves (Fig. 8E). When the oscillation frequency of P_H_ was set to be higher than that of P_V_ (*a*_1_ = 0.36 and *a*_20_ = 0.39), propagated waves initially collided midway, but thereafter the direction was fixed to be from P_H_ to P_V_ (Fig. 8F, top). When the frequencies of P_H_ and P_V_ were closer (*a*_1_ = 0.38 and *a*_20_ = 0.39), the direction of waves was fixed from P_H_ to P_V_, but the period for initial collisions became longer (Fig. 8F, bottom). Importantly, when the oscillation frequency of P_H_ was higher (*a*_1_ = 0.30), the different parameter values of P_V_ (*a*_20_ = 0.36, 0.40, and 0.44) did not affect the wave direction or frequency represented after the period of collisions (Fig. 8G). These results indicated that when the intrinsic frequencies expressed by remotely coupled pacemakers were different, propagated waves were directed from one oscillatory pacemaker with a higher frequency to the other, and unless collisions occurred, the potential to express oscillations of the recessive P cell was totally masked.

### Temporal changes in the parameter in pacemakers sufficiently lead to heart reversals

A previous study based on a different mathematical model proposed that gradual change in the pacemaker firing rate is a possible mechanism for heart reversals (Cain et al., 2020). We also tested a possible mechanism for heart reversals in our model. We increased the parameter “*a*” in P_H_ from the oscillatory mode (from *a*_1_ = 0.37 at a rate of *da*/*dt* = 0.005 during 40 ≤ *t* ≤ 60) and also decreased that in P_V_ from the excitatory mode (from *a*_20_ = 0.44 at a rate of *da*/*dt* = −0.005 during 41 ≤ *t* ≤ 61) (Fig. 9A, upper). In this case, waves propagated from P_H_ to P_V_ as long as these maintained their respective modes (Fig. 9A, upper). However, waves paused as the parameter of P_H_ (*a*_1_) became larger than that of P_V_ (*a*_20_, at *t* = 47 or later), at which point, P_H_ and P_V_ both became excitatory (Fig. 9A, arrowhead). AP waves propagated in the reverse direction from P_V_ to P_H_, after P_V_ was set in the oscillatory mode (Fig. 9A, upper). When the decrease in *a*_20_ started slightly earlier (during 40 ≤ *t* ≤ 60), the duration of the pause was shortened (Fig. 9A, lower). This simulation pattern corresponded well to the reversal with a pause observed in the actual isolated *Ciona* heart (Fig. 9B).

**Fig. 9.**
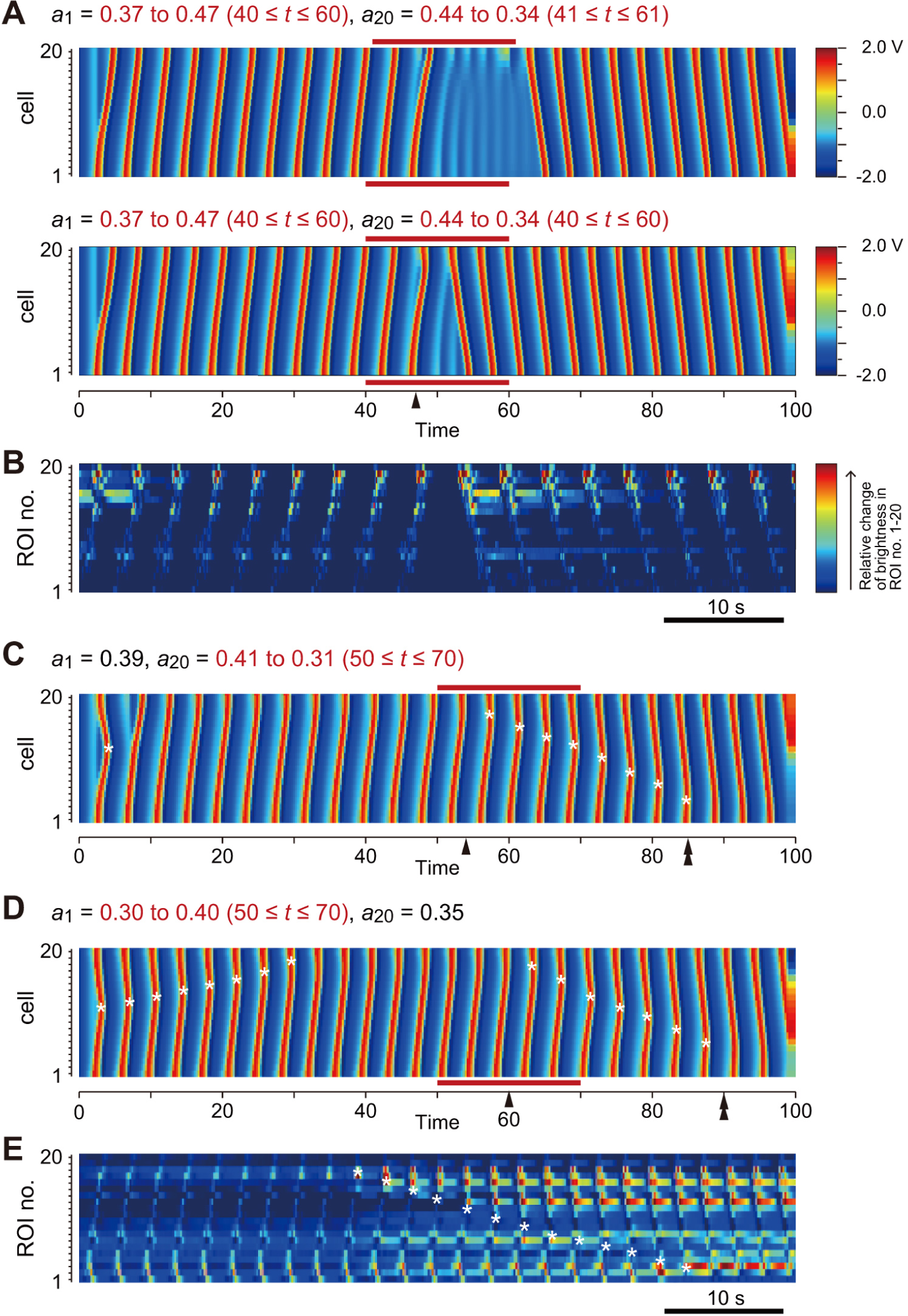
Frequency changes in P cells lead to heart reversals in the model. (A) When we increased *a*_1_ from 0.37 to 0.47 during 40 ≤ *t* ≤ 60 and decreased *a*_20_ from 0.44 to 0.34 during 41 ≤ *t* ≤ 61, a temporal pattern similar to heart reversals with a pause was produced (upper). When a decrease in *a*_20_ was applied during 40 ≤ *t* ≤ 60, the pause was shortened (lower). The arrowhead indicates the time point at which *a*_1_ and *a*_20_ both equaled 0.405. (B) Relative changes in brightness in the twenty ROIs placed along a *Ciona* heart tube representing the propagated waves of pulsations. A heart reversal with a pause is shown as a heat map. (C) When we fixed *a*_1_ = 0.39 and decreased *a*_20_ from 0.41 to 0.31 during 50 ≤ *t* ≤ 70, a temporal pattern similar to heart reversals with collisions was produced. At *t* = 52, P_V_ was set to the oscillatory mode, and *a*_20_ became smaller than *a*_1_ at *t* = 54 or later (arrowhead). Collisions ended around *t* = 85, and the heart reversal was accomplished (double arrowhead). (D) When we increased *a*_1_ from 0.30 to 0.40 during 50 ≤ *t* ≤ 70 and fixed *a*_20_ = 0.35, a heart reversal was again produced. *a*_1_ became larger than *a*_20_ at *t* = 60 or later (arrowhead) and collisions then started. The heart reversal was completed around *t* = 90 (double arrowhead). (E) Relative changes in brightness in the twenty ROIs placed along a *Ciona* heart tube. A heart reversal with collisions is shown. Asterisks indicate the spatiotemporal points of collisions in C-E.

We tested conditions to reproduce reversals with collisions (see Fig. 4B and Fig. S1). We temporally changed the frequency in the simulation by decreasing the parameter in P_V_ (from *a*_20_ = 0.41 at a rate of *da*/*dt* = −0.005 during 50 ≤ *t* ≤ 70 and *da*/*dt* = 0.0 at *t* ≥ 70; oscillatory mode at *t* ≥ 52) while fixing that in P_H_ to the oscillatory mode (*a*_1_ = 0.39) (Fig. 9C). Under these conditions, propagated waves were initially directed from P_H_ to P_V_ because the intrinsic frequency of P_H_ was higher than that of P_V_ (0 ≤ *t* ≤ 54) (Fig. 9C, arrowhead). When the frequency of P_V_ became equal to or higher than that of P_H_ (54 ≤ *t* ≤ 85), waves propagated from both ends and repetitively collided on the way (Fig. 9C). In the final stage, waves selectively travelled from P_V_ to P_H_, the opposite direction (85 < *t*) (Fig. 9C, double arrowhead).

We verified this differential-frequency pacemaker scenario for the reversal with collisions by temporally changing the parameter in P_H_ instead of that in P_V_ [increasing from *a*_1_ = 0.30 (oscillatory mode) at a rate of *da*/*dt* = 0.005 during 50 ≤ *t* ≤ 70, and *da*/*dt* = 0.0 at *t* ≥ 70 (oscillatory mode), and fixed *a*_20_ = 0.35 (oscillatory)] (Fig. 9D). When the frequency of P_V_ became equal to or higher than that of P_H_ (60 ≤ *t*), a series of collisions were observed (60 ≤ *t* ≤ 90) (Fig. 9D). The reversal was completed and waves propagated from P_V_ to P_H_ (90 < *t*) (Fig. 9D, double arrowhead). These patterns of reversals were similar to the actual heart reversal with collisions observed in the isolated *Ciona* heart (Fig. 9E; cf. Fig. 9C,D). Collectively, these results showed that the FitzHugh-Nagumo model applied to linearly aligned cells sufficiently simulated heart reversals with a pause and those with collisions in the *Ciona* heart tube.

## DISCUSSION

We herein demonstrated that (i) the isolated heart of *Ciona* sufficiently exhibited heart reversals, (ii) pulsation frequencies expressed by H- and V-sided pacemakers were changeable, (iii) heart reversals with a pause accompanied preceding decreases in the frequency, whereas reversals with collisions accompanied overcoming of the pulsation frequency in the hitherto dominant end by that in the hitherto recessive end, (iv) H- and V-sided pacemakers possessed properties to express bimodal rhythms, (v) only the 5% or shorter terminal regions of the heart tube continued expressing autonomous pulsation rhythms, whereas other regions did not. Furthermore, we showed using our mathematical model that (vi) the difference between frequencies expressed by the H and V ends determined the direction of propagated waves, (vii) the intrinsic rhythm of the recessive end was totally masked, and (viii) changes in the statuses (excitatory or oscillatory) and intrinsic rhythms of the terminal pacemakers could lead to heart reversals.

We herein propose the following mechanism for heart reversals in *Ciona*: (I) The heart tube has two remotely coupled pacemakers at each end. (II) These pacemakers express bimodal rhythms, namely their rhythms are periodically accelerated and decelerated or occasionally enter the excitatory mode and return to the oscillatory mode. (III) Based on the differential-frequency pacemaker scenario, pulsation waves are propagated from one pacemaker with an intrinsic status expressing a higher frequency of oscillations to the other, and the direction intermittently reverses according to independent changes in the rhythms expressed by the pacemakers (see also Cain et al., 2020).

### The *Ciona* heart tube has two independent pacemakers in its termini

The pacemakers of the *Ciona* heart tube are considered to reside in terminal regions based on pulsations originating from its termini. However, this is controversial because of experimental findings showing that every fragment taken from any part of the heart tube exhibited autonomous beating (see Krijgsman, 1956; Anderson, 1968; Kriebel, 1968b; Goodbody, 1974). We herein demonstrated that most of the middle fragments excluding the < 5% terminal region, initially exhibited autonomous beating, but all ceased beating within 30 min of fragmentation. On the other hand, fragments including either end continued to express rhythmic pulsations. We concluded that the heart tube fragments prepared in previous studies may have been examined shortly after divisions, and also that the true pacemakers of the *Ciona* heart only reside in the terminal regions.

We repeatedly observed marked elevations in the activity of the heart or its fragments for tens of minutes to a few hours after manipulations. This effect was prominent on the H side. The heart directed pulsation waves from the H end to the V end immediately after its isolation (see Fig. 2), the appearance of bimodal rhythms in the bisected hearts was later on the H side (Fig. 6), more specimens of H-sided middle quarters showed beating after the division, and the pulsation frequencies of H terminal fragments were higher than those of V terminal fragments (Fig. 7). Since the objects were activated in all the cases, like the effect to decrease “*a*” in the FitzHugh-Nagumo model depicted above, we considered that these activations to reflect a common artefactual effect through experimental manipulations, e.g., derived from physical damage to tissues. This artefactual effect, however, appeared to at least partially decrease; e.g., beating frequencies were restored to a moderate level, heart reversals were also revived in a few hours, and so on. The concept of a reversible effect temporarily changing working cardiomyocytes into a pacemaker is intriguing because it implies that pacemakers and working myocytes in the *Ciona* heart tube share basic properties and are not markedly different from each other, just as a slight difference in the parameter “*a*” in our model.

### Bimodal rhythms expressed by terminal pacemakers

The presence of pacemakers in terminal regions does not sufficiently explain the mechanism for heart reversals by itself. Our mathematical model showed that propagated AP waves were directed from the pacemaker in the oscillatory mode with a lower parameter “*a*” value to the other, and the intrinsic rhythm of the recessive P cell was totally masked. This hierarchical relationship of terminal pacemakers, which we herein referred to as a differential-frequency pacemaker scenario, constitutes a sufficient mechanism for heart reversals, under the condition that the rhythms of terminal pacemakers are changeable (see also Cain et al., 2020).

We demonstrated that fragments including either end exhibited bimodal rhythms. Previous studies reported that heart tube fragments showed beating cycle changes (Kriebel, 1968a, 1968b; Anderson, 1968). Anderson (1968) indicated that the V-sided halves of the *Ciona* heart tube exhibited a regular cycle of accelerations and decelerations of the beating rate and discussed its relationship with heart reversals. We also propose that this ability to express bimodal rhythms allows the *Ciona* heart pacemakers to lead heart reversals.

The mechanisms underlying bimodal rhythms remain unclear. The repetitive “bursting” of APs in various types of neurons and endocrine cells (Plant and Kim, 1976; Chay and Rinzel, 1985) and other phenomena, including “Cheyne-Stokes respiration”, in which the periods for shallow and deep breathing repetitively alternate (e.g. Leung and Bradley, 2003), also represent biological bimodal rhythms. The latter cases are often observed in patients with some degree of heart failure, suggesting that a change in the physiological status induces a transition from a regular rhythm to bimodal rhythms. We do not necessarily support the previous proposal of “fatigue” in the dominant pacemaker eliciting heart reversals (Krijgsman, 1956) since we observed robust pulsations and reversals in hearts 1 day after their isolation. However, it seems reasonable to hypothesize the gradual accumulation of some metabolites in the P cell(s) during its dominant (or recessive) periods, and the modulation of the pacemaking mechanism by the metabolite(s) as second messenger(s).

### Pulsations in the heart within or outside the body

Whether the pacemaking mechanism for the heartbeat resides in the heart itself or in the central nervous system has been an area of interest in invertebrate biology (e.g. Kodirov et al., 2018). We and previous studies showed long-lasting pulsations in the isolated heart of *Ciona*, and, thus, a central pattern generator is not required for the heartbeats in this animal. However, we do not deny the possible presence of some neural inputs into the heart in *Ciona*. In the present study, the pulsation frequency during H-dominant periods decreased, and, accordingly, the duration of the H-dominant period was irreversibly shortened in isolated hearts. A cholinergic input into the heart was previously reported in the colonial ascidian *Botryllus schlosseri* (Burighel et al., 2001). Some factors inside the body, including autonomic nervous controls, may support the dominance of the pacemaker in the H end. The presence of the pericardial wall may also be critical, since the excision of the pericardial wall led to irreversible increases in the incidence of collisions. A suitable humoral environment and/or physical pressure may also contribute to ensuring that the H-end pacemaker is relatively dominant (Waldrop and Miller, 2015).

A recent study also informed us that a network of cells expressing the peptide proprotein convertase 2 gene ortholog, one of the reliable markers for peptide-producing endocrine/neuroendocrine/neuronal cells, was closely associated with the H and V ends and the raphe (Osugi et al., 2020). This network complicates the issue of whether autonomous pulsations in the isolated *Ciona* heart are myogenic or neurogenic. Another study showing that the expression of an HCN channel, a key factor for spontaneous AP generation, in the heart tube cells of *Botryllus* supported the heartbeat rhythm being myogenic also in the ascidian heart (Hellbach et al., 2011).

### Hearts of chordates and possible causes for heart reversals in ascidians

The vertebrate heart has at least two major populations of pacemaker cells that express independent pulsation rhythms: the sinoatrial node (SAN) and the atrioventricular node (AVN) (e.g. van Weerd and Christoffels, 2016). Although it is premature to speculate whether these systems of two pacemakers in *Ciona* and vertebrate hearts are homologous (Burkhard et al., 2017; Cain et al., 2020), the autonomous rhythm of the isolated SAN was shown to be higher than that of the isolated AVN, with the former representing the dominant pacemaker in the vertebrate heart (e.g. Gaskell, 1883).

Drastic changes in the heartbeat rhythm and blood pressure are inadequate for vertebrate life. However, since ascidians are sessile and their metabolism rate is markedly low, they are more likely to be tolerant of irregular pacemakers in their hearts. Reversely, these irregular pacemakers may be advantageous for this animal. The open circulatory system in ascidians includes narrow and complex channels, such as sinuses in the branchial basket, which get crumpled during “squirting”. Heart reversals may relieve the clogging of blood cells and deliver fresh body fluids to otherwise congested sites, as originally proposed (Kuhl and van Hasselt, 1822; for review, Krijgsman, 1956). In order to solve the functional purpose of heart reversals, their temporal pattern is expected to be experimentally altered to examine its effects on viability, growth, or reproductive success. The present study provides basic implications to approach these unsolved issues; e.g., the ultimate causes of ascidian heart reversals and the phylogenetic origin of the chordate heart architecture.

## ACKNOWLEDGMENTS

We thank Drs. Hiromi Kakizaki, Satoe Aratake, Megumi Kotsuka, Akihiro Yoshikawa, Makoto Nozawa, and Manabu Yoshida (Misaki Marine Biological Station, The University of Tokyo, Japan), Drs. Reiko Yoshida, Chikako Imaizumi, and Yutaka Satou (Laboratory of Developmental Genomics, Department of Zoology, Graduate School of Science, Kyoto University, Japan), and other staff who distribute *Ciona* under the National BioResource Project, AMED, Japan. We also thank Drs. Yasushi Okamura, Fumihito Ono, Koichi Nakajo, and Yasunori Sasakura for their encouragement.

## DATA AVAILABILITY

All the experimental data are found in the article and the supplements. The source codes are available upon request.

